# Microbial communities across a hillslope-riparian transect shaped by proximity to the stream, groundwater table, and weathered bedrock

**DOI:** 10.1101/423368

**Authors:** Adi Lavy, David Geller McGrath, Paula B. Matheus Carnevali, Jiamin Wan, Wenming Dong, Tetsu Tokunaga, Brian C. Thomas, Kenneth H. Williams, Susan Hubbard, Jillian F. Banfield

## Abstract

Watersheds are important suppliers of freshwater for human societies. Within mountainous watersheds, microbial communities impact water chemistry and element fluxes as water from precipitation events discharges through soils and underlying weathered rock, yet there is limited information regarding the structure and function of these communities. Within the East River, CO watershed, we conducted a depth-resolved, hillslope to riparian zone transect study to identify factors that control how microorganisms are distributed and their functions. Metagenomic and geochemical analyses indicate that distance from the East River and proximity to groundwater and underlying weathered shale strongly impact microbial community structure and metabolic potential. Riparian zone microbial communities are compositionally distinct from all hillslope communities. Bacteria from phyla lacking isolated representatives consistently increase in abundance with increasing depth, but only in the riparian zone saturated sediments did we find Candidate Phyla Radiation bacteria. Riparian zone microbial communities are functionally differentiated from hillslope communities based on their capacities for carbon and nitrogen fixation and sulfate reduction. Selenium reduction is prominent at depth in weathered shale and saturated riparian zone sediments. We anticipate that the drivers of community composition and metabolic potential identified throughout the studied transect will predict patterns across the larger watershed hillslope system.

## Introduction

Soil microbial communities impact our environment by driving biogeochemical cycles from centimeter to global scales (Schimel and Schaeffer, 2012; Rousk and Bengtson, 2014). They expedite rock weathering (Krumbein, 1988; Gorbushina, 2007) recycle organic material in the subsurface, and facilitate the growth of vegetation by altering the availability of nutrients in the soil (Wardle *et al.*, 2004). These changes impact soil nutritional status and productivity and plant survival and biotic interactions. It is not surprising then, that microbial communities are considered ecosystem engineers (Viles, 2012).

In recent years, mountain watersheds have received attention because of their importance for supplying most of the world’s fresh water (Viviroli *et al.*, 2003) and because of their contributions to subsurface carbon storage (Hagedorn *et al.*, 2010; Chang *et al.*, 2014; Wan *et al.*, 2018) These environments are comprised of a complex system of components, such as forests and meadows, floodplains, and glaciers. In turn, each of these accommodates various habitats including soil, bare rock, permafrost, and snow. Development of a predictive understanding of the behavior of such a heterogeneous and interconnected set of ecosystem compartments is an extremely complicated undertaking. Employing a scale-adaptive approach in which different ecosystem compartments are considered as “systems within systems” could assist in disentangling the processes that shape overall mountain ecosystem function (Levin, 1992; Hubbard *et al.*, 2018). A first step toward such a goal is to investigate structure and functioning within individual montane ecosystem compartments to provide a basis for future comparative studies and modeling efforts. In the long term, the “systems within systems” approach may better enable predictions accompanying natural or anthropogenic environmental perturbations.

Hillslope and floodplain compartments host the majority of soils in alpine and subalpine mountain ecosystems, and biogeochemical processes that occur there impact downstream ecosystems. Runoff and groundwater transport solutes along the elevation gradient and into aquifers, rivers and lakes. Soils ay hillslopes and floodplains, and in general, harbor considerable microbial diversity (Rime *et al.*, 2014; Frey *et al.*, 2016; Donhauser and Frey, 2018). Most studies of microbial communities in mountainous soils have been concerned with the microbial community structure across different climate zones on the mountain slopes (Djukic *et al.*, 2010; Zhang *et al.*, 2013; Xu *et al.*, 2014; Klimek *et al.*, 2015; Bardelli *et al.*, 2017). However, most work has focused only on shallow soil, down to 20 cm (Zhang *et al.*, 2013; Yuan *et al.*, 2014; Bardelli *et al.*, 2017) and sometimes only the top 5 cm (Singh *et al.*, 2014). The shallow layer of soil is profoundly affected by low temperatures that frequently drop below 0 °C and snow cover that crucially limits biological, chemical and physical processes, and thus microbial life (Zumsteg *et al.*, 2013). In contrast, the deeper soils and weathered rock in mountain ecosystems have been little studied. While affected by events taking place in shallow layers, the microbial communities there are probably also strongly influenced by moisture gradients and the geochemistry of the underlying bedrock.

The East River headwaters catchment is a mountainous, high-elevation watershed, dominated by the Cretaceous Mancos Shale Formation, with carbonate and pyrite contents of roughly 20% and 1%, respectively (Morrison *et al.*, 2012). The watershed has a mean annual temperature of ∼0 °C, with average minimum and maximum temperatures of −9.2 °C and 9.8 °C, respectively. The watershed receives ∼600 mm of precipitation per year, the bulk of which falls as snow, and is representative of many other headwaters systems within the upper Colorado River Basin (Pribulick *et al.*, 2016; Hubbard *et al.*, 2018).

The present research focused on a lower montane hillslope through floodplain transect located within the East River, CO watershed of the Lawrence Berkeley National Laboratory-led Watershed Function Project. The intensive site that is the focus of this study is referred to as PLM (Pump House Lower Montane). This scale-adaptive investigation of the Watershed Function Project at the East River watershed explores how mountainous watersheds retain and release downgradient water, nutrients, carbon, and metals (Hubbard et al., 2018). Here, we address the composition, diversity, potential metabolism of microbial communities, and the potential overlap in community composition among sites along an altitudinal transect at different soil-weathered rock depths and examine finer-scale variation within the hillslope system and its intersection with the floodplain riparian zone.

## Methods

### Site description and sample collection

The PLM intensive study site is located on the north-east facing slope of the East River valley near Crested Butte, Colorado, USA (38°55’12.56’N, 106°56’55.39”W) (Fig. 1 and Figure S1). Exact locations were determined at an accuracy of 0.5 meters with a Trimble Geo 7X GPS (Sunnyvale, California, USA). All samples were collected during three days in September 2016 from meadow sites before any intensive research activities were performed. The ground surface at each site was cleared of vegetation with a hand trawler prior to sampling. Samples were collected with a manual corer lined with 7.6 cm tall and 15.2 cm diameter bleached sterile plastic liners. Five soil profile sampling sites, abbreviated PLM0, PLM1, PLM2, PLM3, and PLM4 were chosen along a 230 m hillslope transect. The profiles terminated at depth in the unsaturated zone, with the exception of PLM4, which extended below the water table. The base of PLM3 and PLM3 profiles are located near or within the weathered Mancos Shale bedrock, while the base of PLM0 was located >1 m above the weathered bedrock. PLM0 is at the top of the hill and PLM4 on the East River floodplain, 2788 m and 2743 m above sea level, respectively (Fig. 1). The soil in between sampling depths was removed with an auger. An additional site, PLM6, was sampled by drilling and provided access to weathered shale. Samples at PLM6 were taken from a split-spoon, dry drilled core.

**Figure 1:**
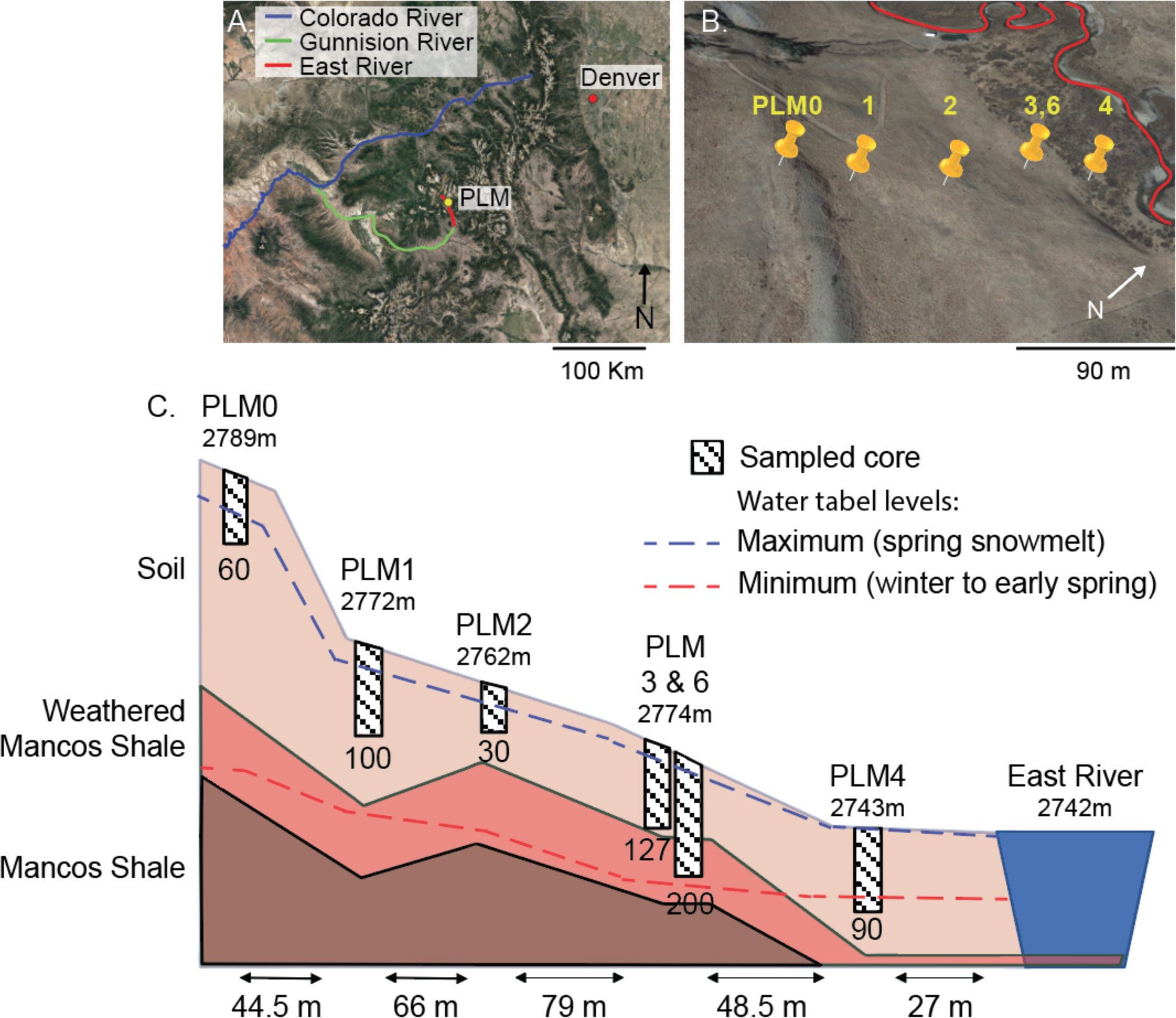
East River Watershed hillslope-riparian zone transect sampling sites. **(A)** The location of East River PLM intensive study site. (B) Five PLM sites are located on a hillslope transect, where PLM0 is at the highest point of the transect and PLM4 is located in the floodplain. (C) Schematic representation of the sampling sites. Elevation of the surface, given in meters above sea level, appears below the name of sampling site. Maximum depth at each sampling site is specified below the depiction of the sampled core. Horizontal distances between sites are given at the bottom of the illustration. Maximum and minimum water levels are depicted by dashed blue and red lines, respectively. PLM6 site was initially drilled for the purpose of another study, 5 m apart and at the same elevation as PLM3. A full view of the East River watershed is given in Figure S1.

Immediately after extracting samples, they were placed in sterile Whirl-Pak bags and manually homogenized. Aliquots of 5 g of soil were placed in 10 ml of LifeGuard Soil Preservation Solution (Qiagen, Netherlands). Care was taken to avoid roots and small rocks. Samples were placed in a chilled cooler until processing at the Rocky Mountain Biological Laboratory (RMBL) later that day. In the laboratory, roots and small rocks were removed from sampling bags and three 10 g subsamples were weighted from each sample and placed in a −80 °C freezer. Samples were shipped overnight on dry ice to University of California, Berkeley for DNA and RNA extractions.

Particle size analyses of samples from PLM0, PLM1 and PLM3 were conducted according standard methods (Gee and Or, 2002). Geochemical measurements were made at the Earth and Environmental Sciences department’s Aqueous Geochemistry Lab. Water soluble cation-anion composition was measured by water extraction (1:1 soil:DIW mass ratio) and ICP-MS. Total inorganic carbon (TIC) and total organic carbon (TOC) in soil samples were determined using a Shimadzu TOC-VCSH total inorganic and organic carbon analyzer combined with a solid sample combustion unit of SSM-5000A (Shimadzu, Kyoto, Japan). Total nitrogen (TN) was analyzed using a Shimadzu Total Nitrogen Module (TNM-1) combined with the TOC-VCSH analyze. Geochemical measurements for samples taken at PLM6, the nitrate concentration for the sample from PLM0 30 cm, and sulfate concentrations for samples PLM0 40 cm, PLM1 60 cm, PLM1 90 cm, PLM2 5cm and PLM2 30cm were not available.

### DNA extraction and sequencing

DNA was extracted from 10 g of soil with DNeasy PowerMax Soil Kit (Qiagen, Netherlands) in two batches of 5 g each, following the manufacturer’s protocol with the following modifications: soil was vortexed at maximum speed for an additional 3 min in the sodium dodecyl sulfate reagent, and then incubated for 30 min at 60 °C, with intermittent shaking in place of extended bead beating as established by Hug et al. (2015). For DNA precipitation, sodium acetate (1:10 v/v) and isopropanol (1:1 v/v) were added and samples were incubated overnight (−20 °C). Following incubation, DNA was pelleted by centrifugation (15300 g, 15 min, 4 °C), washed with cold ethanol and suspended in ddH_2_O. DNA was further cleaned with DNeasy PowerClean Pro Clean Up Kit (Qiagen, Netherlands) following the manufacturer’s protocol. DNA extractions from both batches were combined during the cleaning step.

DNA was also co-extracted with RNA from 5 g of soil using RNeasy PowerSoil Total RNA Kit (Qiagen, Netherlands) and Phenol:Chloroform:Isoamyl Alcohol 25:24:1 saturated with 10 mM Tris (final pH 8.0) and 1 mM EDTA. RNeasy PowerSoil DNA Elution Kit (Qiagen, Netherlands) was used to collect DNA which was further cleaned using DNeasy PowerClean Pro Clean Up Kit (Qiagen, Netherlands). The co-extraction and cleaning steps were conducted according to the manufacturer’s protocol.

Metagenomic libraries were prepared at the Joint Genome Institute (JGI) after validating concentrations and DNA integrity using Qubit (Thermo Fisher Scientific) and gel electrophoresis, respectively. Libraries were prepared using NEB’s Ultra DNA Library Prep kit (New England Biolabs, MA) for Illumina with AmpureXP bead selection aimed to give fragments of 500 bp according to the manufacturer’s protocol. The library was sequenced at JGI using an Illumina Hiseq 2500, resulting in paired-end, 150 base-pair (bp) sequences.

### Bioinformatic analysis

Raw reads processing followed protocols described elsewhere (Hernsdorf *et al.*, 2017). Briefly, reads were trimmed based on quality scores with Sickle (Joshi and Fass, 2011) and assembly was accomplished with IDBA-UD v1.1.1 (Peng *et al.*, 2012) using kmer size range of 40-140. Only assembled scaffolds with > 1 Kbp were included in downstream analysis. Open reading frames were identified by Prodigal v2.6.3 (Hyatt *et al.*, 2010) using the metagenomic setting.

Microbial community structure was assessed according to the abundance of ribosomal protein S3 (rpS3) genes by modifying the method described by Anantharaman et al. (2016). Archaeal, eukaryotic and bacterial rpS3 protein sequences were identified using Hidden Markov Models (HMM) (Finn *et al.*, 2015). Ten rpS3 reference sequences which compose TIGRFam’s TIG01009 model were added to the protein sequences that were identified by HMMs and aligned with MAFFT (Katoh and Standley, 2013). Positions within the alignment with >95% gaps were removed, leaving 206 amino acids in the longest, non-reference sequence. Sequences that had less than 103 non-gap positions (50% of overall non-gap positions) were removed from the analysis. This step ensured that only positions that are truly related to the sequence of rpS3 were included in downstream analysis.

The amino acid sequences were clustered with the cluster_fast algorithm from usearch software (Edgar, 2010) at a 99% similarity threshold, and the following settings: query_cov=1, target_cov=0.5, and both max_accept and max_reject set to 0. Scaffolds of DNA sequences that matched the clusters’ open reading frames were retrieved from the metagenomes. Average coverage was used as a proxy for relative abundance of different sequence types. In this analysis, the scaffolds were trimmed to include 2 Kbp flanking the rpS3 gene. If the scaffold did not span 2 Kbp on either side, then the entire scaffold was kept, with a minimal length of 1 Kbp. The relative abundance of each trimmed scaffold was determined by mapping the reads from each sample to each trimmed scaffold with bowtie2 (Langmead and Salzberg, 2012). The average coverage and breadth of coverage of each scaffold in each sample was then calculated (Olm *et al.*, 2017). Each scaffold is considered to be present in at least one sample (at minimum, the sample from which it was originally assembled) but could be falsely identified in other samples due to a low breadth cutoff (i.e., false positive). Therefore, we implemented a breadth cutoff of 0.72 based on iterating breadth cutoffs of 0.1 to 1, to find the lowest breadth cutoff that would retain the same number of clusters as went into the analysis.

Genes involved in carbon, nitrogen and sulfur metabolism were identified using 114 previously published HMM models (Anantharaman *et al.*, 2016) (Table S1). Three genes encoding for enzymes taking part in selenium metabolism were also sought for using TIGRFAM HMM models (Table S1). Additionally, *srdA* which encodes for a membrane bound catalytic subunit of selenate reductase, was detected with a custom HMM model. The model was constructed by aligning 20 amino-acid sequences, 934-1222 aa long, determined to be included in the *srdA* specific clade (Harel *et al.*, 2016). All matches from HMM search for *srdA* were aligned and a threshold was decided upon according to their clustering in a phylogenetic tree. The abundance of each gene was determined by mapping the reads from each sample to each gene as was done for rpS3 gene. The same breadth cutoff was used as before. Cumulative abundance was transformed with regularized logarithm (RLD) following DESeq2 analysis (Love *et al.*, 2014), and only HMMs with transformed cumulative abundance >1 were considered in further analysis.

### Taxonomy and phylogeny

The longest amino acid sequence from each rpS3 protein sequence cluster was selected as a representative and was compared to a database of rpS3 protein sequences (Hug *et al.*, 2016) using the ublast function in usearch (Edgar, 2010). Results were filtered to include only the top hits with e-values lower than 1e-5. While each cluster roughly correlates with a species, not all clusters could be taxonomically identified to that level. Therefore, further investigation relied on phylogenetic distance which enables a high-resolution analysis. A phylogenetic tree was created by aligning only the representative amino acid sequences using MAFFT with an automated strategy (Katoh and Standley, 2013) and trimming non-informative positions. A Maximum-Likelihood tree was constructed on CIPRES (Miller *et al.*, 2010) with RAxML (Stamatakis, 2014), using the LG substitution model and bootstrapping, allowing the software to halt bootstrapping once it reached a consensus. The Eukaryote domain branch was set as root, and the tree was manually inspected for errors. The phylogenetic tree along with rpS3 gene abundance heatmap were visualized with iTol v4.2.3 (Letunic and Bork, 2016).

#### Statistics

Statistical analysis was conduct in R v3.4.3 (R Development Core Team, 2012) and Rstudio v1.1.423 (Rstudio Team, 2015). Abundance plots, ordinations and Unifrac calculations were conducted with Phyloseq v1.22.3 (McMurdie and Holmes, 2013). The abundance of each rpS3 cluster was corrected for uneven sequencing depth across samples by multiplying the coverage value for each sample by a factor calculated as the ratio of the number of bp in the largest sample divided by the number of bp in that sample.

Factor selection of soil chemistry was carried with vif.cca and bidirectional ordiR2step functions from Vegan v2.4.6 (Oksanen *et al.*, 2018). The best set of explaining factors was identified with bio.env function, with a Euclidean distance method, Bray-Curtis matrix and evaluated with Spearman correlation. Maps were retrieved from Google maps database using Google Earth v7.3.2.

#### Data availability

All reads are available through the NCBI Short Reads Archive. Accession number for each sample is provided in supplementary Table S2.

## Results

For the hillslope samples analyzed, the soils are loamy to silty-loam and the soil texture is more similar in deeper samples compared to shallow samples (Fig. S2 and Table S3). For example, shallow samples from PLM0 and PLM1 have higher sand content than downslope PLM3 samples, which have higher content of clay and silt, potentially as a result of downslope fining of transported sediments. Soil moisture increases with proximity to the East River, but decreases with depth (Fig. S2 and Table S4). An exception to this is at the floodplain, where moisture increases close to the water table (72 cm below the ground surface at the time of sampling). The hillslope meadow is dotted with smooth brome (*Bromus inermis*) and lupines (*Lupine* sp.); however, neither occurred within a 50 cm radius of the sampling sites (qualitative assessment on site). In contrast, the floodplain is dominated by willows and sedges that are not present on the hillslope. Gopher activity increases downslope, but does not occur at the floodplain location (Wendy Brown, personal communication).

Assembling reads from 41 samples, comprising 610 Gbp of sequence data, resulted in 6.5 million scaffolds longer than 1 Kbp (Table S1). Encoded on these, 3536 rpS3 amino acid sequences were identified and clustered into 1660 clusters (at 99% identity), representing 37 microbial phyla. In general, the microbial communities are dominated by bacteria (relative abundance 0.95 ±0.03 sd). The most abundant phyla across all samples are Acidobacteria, Actinobacteria, Chloroflexi and Proteobacteria, but their relative abundance varies considerably across samples and depths (Fig. 2). Species of Verrucomicrobia and to a lesser extent also Gemmatimonadetes are more abundant at sites high on the hillslope (i.e., PLM0, PLM1, and PLM2) compared to their abundance at PLM3 and the floodplain site PLM4.

**Figure 2:**
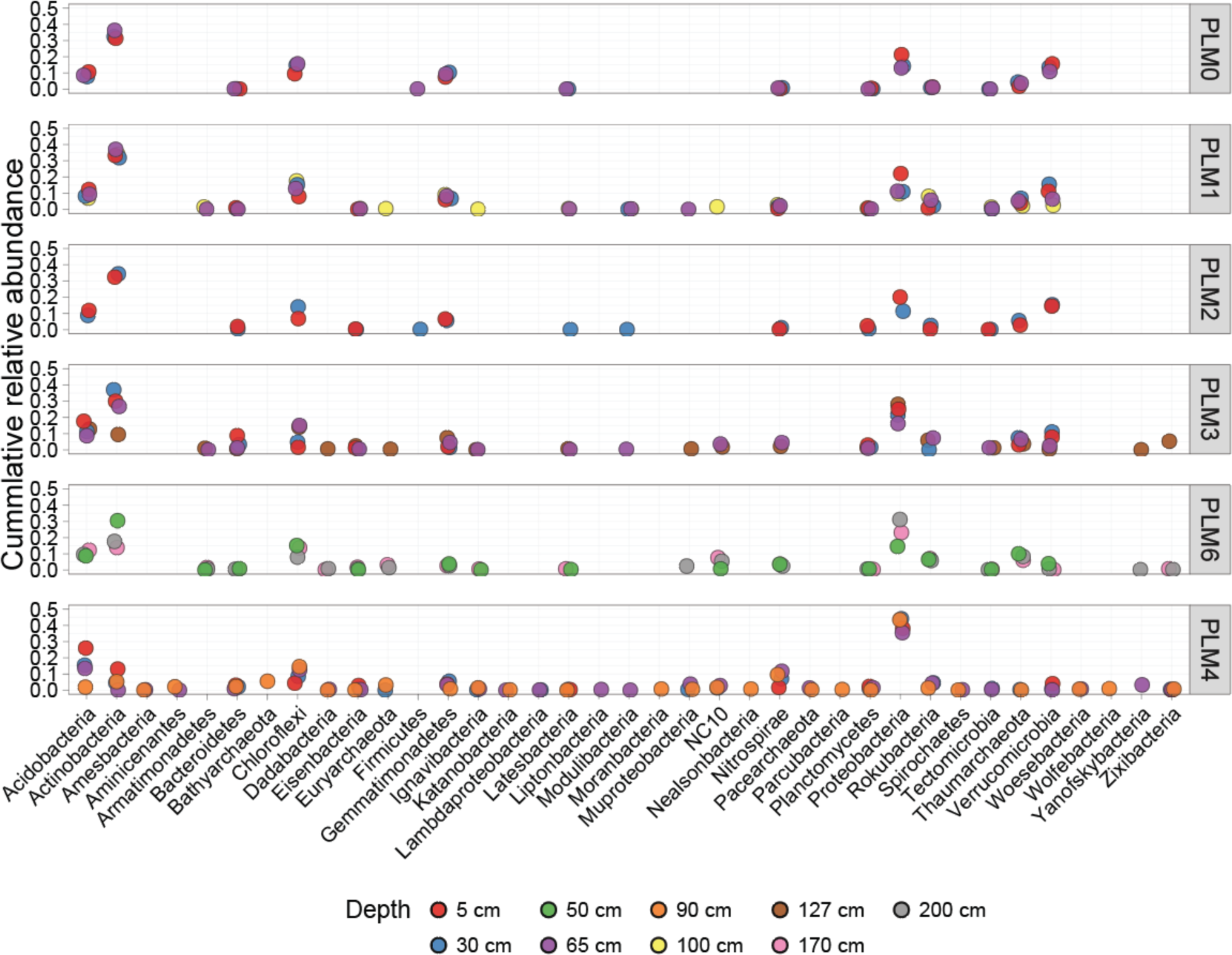
Cumulative relative abundance of phyla. Colors represent depths from which phyla were identified. Results show that Actinobacteria decrease in abundance with increasing depth and proximity to the floodplain site PLM4; Verrucomicrobia show the opposite pattern.

Proteobacteria species comprise 22.7% (±10.8 sd) of all microbial abundance. This dominance increases systematically with distance down the hillslope, largely irrespective of the sampling depth (Fig. 2 and Figure 3A). Gammaproteobacteria species are almost undetectable in communities higher on the hillslope, whereas alphaproteobacterial species are prevalent at all sites (Fig. 3A). Deltaproteobacteria species increase in abundance with increasing proximity to the floodplain and also with increasing proximity to the water table, with the highest representation observed in samples from below the water table. Distinct Deltaproteobacteria species are found in samples close to the water table (*Desulfobacca acetoxidans* in clade 1, and *Geobacter* spp. and *Desulfuromonas* sp. in clades 3 and 4, see Figure S3). Some distinct species (clade 2 in Figure S3) occur only below the water table (Syntrophaceae, Figure S3, clade 2). Thaumarchaeota related to *Nitrososphaera* sp. are the dominant archaea at every location other than at the floodplain (Fig. 3B). At the floodplain site (PLM4), Pacearchaeota are present in soil samples close to, although above the water table whereas Bathyarchaeota and Euryarchaeota are present in samples below the water table.

**Figure 3:**
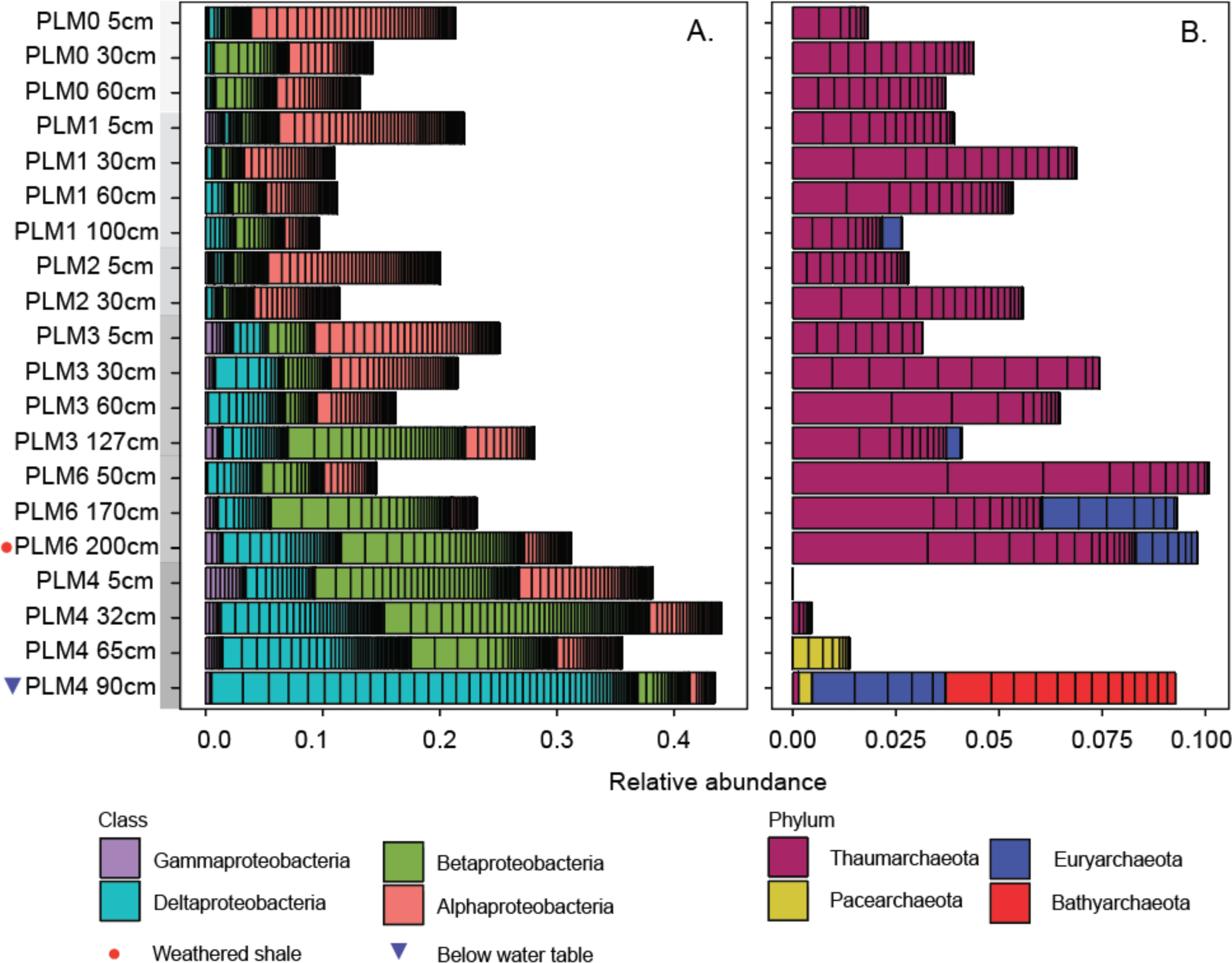
Relative abundances of proteobacterial classes (A) and archaeal phyla (B) clusters across the sampling sites. Within bars of the same color, black lines separate distinct organisms. Samples are ordered from the top to the bottom of the hillslope transect. Within each site, samples are ordered by depth.

Out of the 37 microbial phyla that were identified, 20 are candidate phyla (CP) (i.e., phyla that lack an isolated representative). Of the CP, eight are part of the Candidate Phyla Radiation (CPR) (Fig. 4). Members of CP are present at all sites along the hillslope transect, but they are generally more abundant in deeper samples at each site (Fig. 4A). Interestingly, CPR bacteria are almost exclusively found at the floodplain site and only just above and just below the water table (Fig. 4B). Although sampling sites above and below the water table are close spatially and may experience similar conditions when groundwater level fluctuate, they harbor bacteria from completely different CPR phyla.

**Figure 4:**
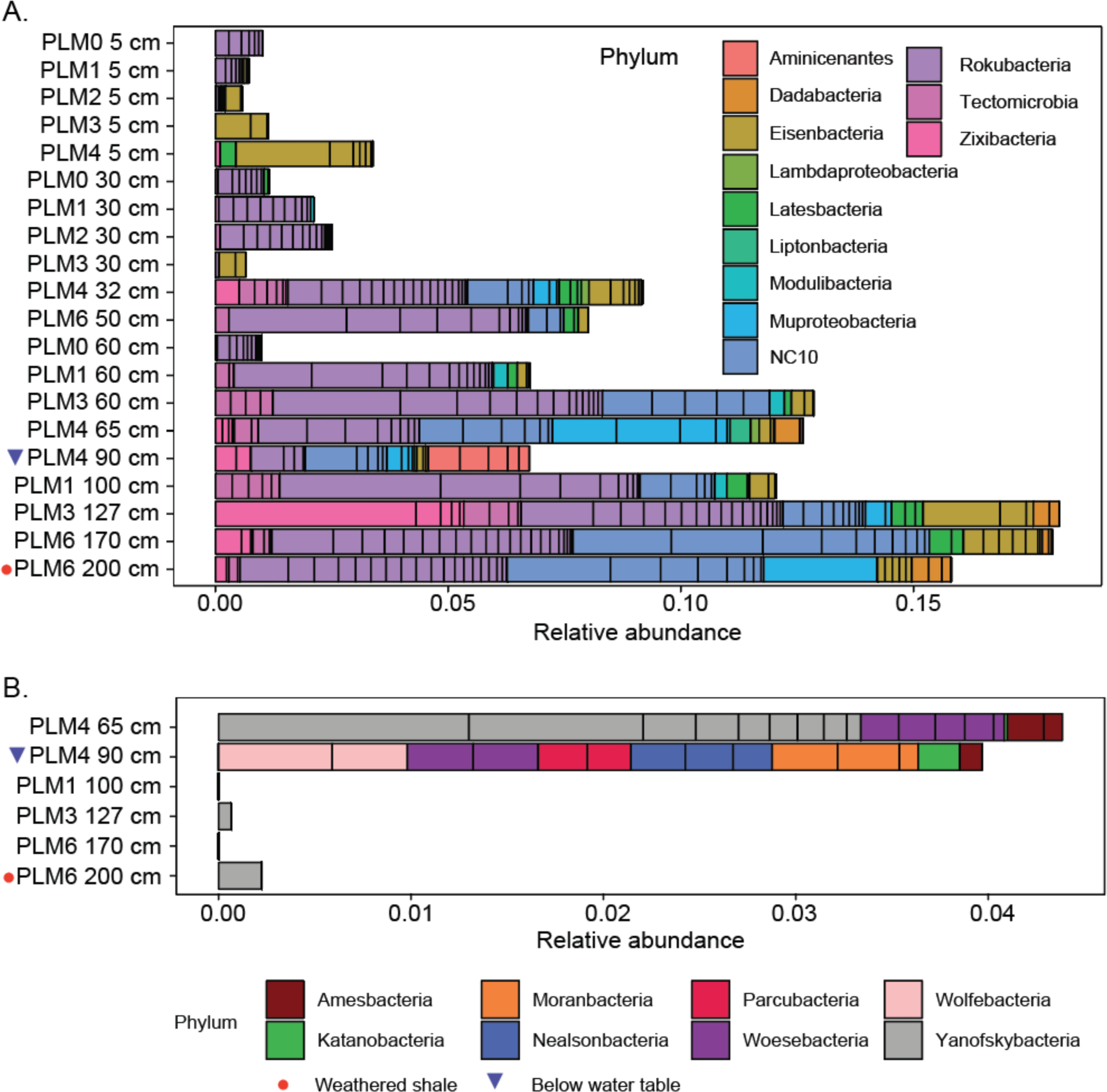
Abundances of Candidate phyla and Candidate Phyla Radiation clusters at hillslope sites. (A) Abundance of bacteria from Candidate phyla other than CPR phyla. (B) Abundance of bacteria from CPR phyla. Samples are ordered by depth and within any specific depth, from top to bottom of the hillslope transect.

We investigated how distance from groundwater and weathered shale impact microbial community structure. Unweighted Unifrac based PCoA ordination reveals that soils sampled at depths of 5 cm and 30 cm from all sites except at the floodplain (PLM4) group together (Fig. 5A). However, the weighted Unifrac PCoA analysis (considering organism abundances) differentiates these 5 cm from 30 cm soil samples and also separates samples from PLM4 from above and below the water table (Fig. 5B). Unweighted Unifrac based PCoA indicates that communities in weathered shale sampled at PLM6 and proximity to weathered shale from PLM3 have similar microbial community compositions (Fig. 5A), but this similarity decreases when organism abundances are considered (Fig. 5B). Thus, for soils that contain similar types of organisms, sampling depth and proximity to weathered rock shift organism abundance relative levels. Overall, distance from groundwater and weathered shale seem to be dominant factors in determining the microbial community structure across the hillslope.

**Figure 5:**
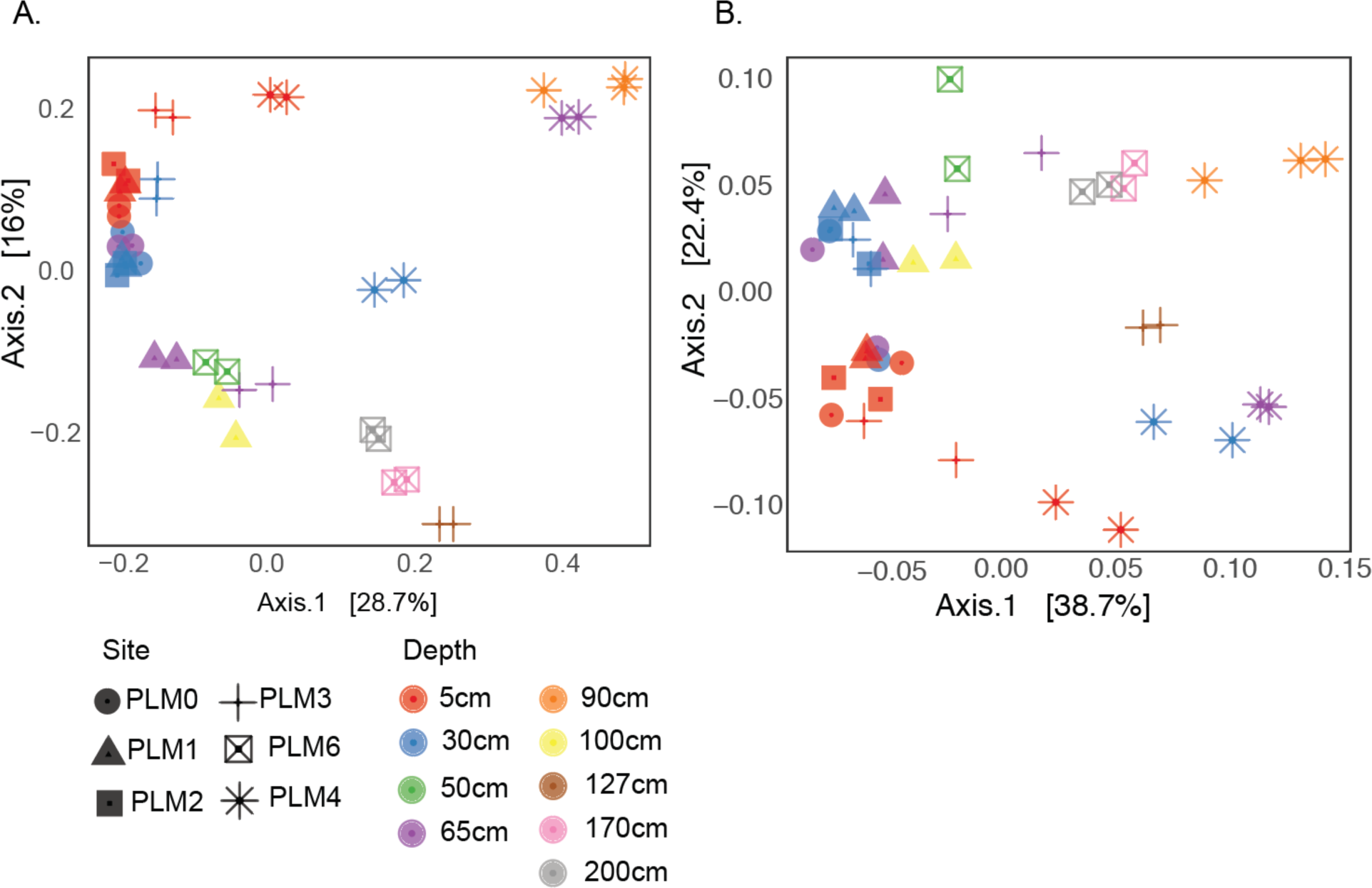
Samples cluster based on proximity to weathered shale and groundwater-saturated soil. (A) Principal Component Analysis (PCoA) based on unweighted UniFrac distance computed using Maximum-Likelihood phylogenetic tree. (B) PCoA based on weighted UniFrac distances computed using Maximum-Likelihood phylogenetic tree and abundance of each taxon. Numerical values in parentheses next to names of axis shows the percentage of variability that can be explained by the location along that axis.

Fourteen geochemical factors were assessed in order to elucidate the factors that shape community structure in the soil profile sites. The combination of soil moisture and concentrations of Se, Mo, and Zn were correlated to microbial community structure (rho=0.809) (Fig. 6). As single factors, Se (adjusted r^2^=0.124, p-value=0.01), Zn (adjusted r^2^=0.212, p-value=0.006) and soil moisture (adjusted r^2^=0.279, p-value=0.022) show significant correlations. Selenium had the highest concentration in samples taken above the water table, (PLM4 65 cm, 8.119 ± 0.235 ppb) whereas sulfate and molybdenum concentrations were the highest in deeper samples (PLM4 90 cm, 6.68 mg/L and 12.69 ppb, respectively) (Fig. S3). Zinc concentration was the highest in samples closest to weathered shale (PLM3 127 cm, 95.694 ± 0.915 ppb), which also had the lowest acidity (pH=7.98).

**Figure 6:**
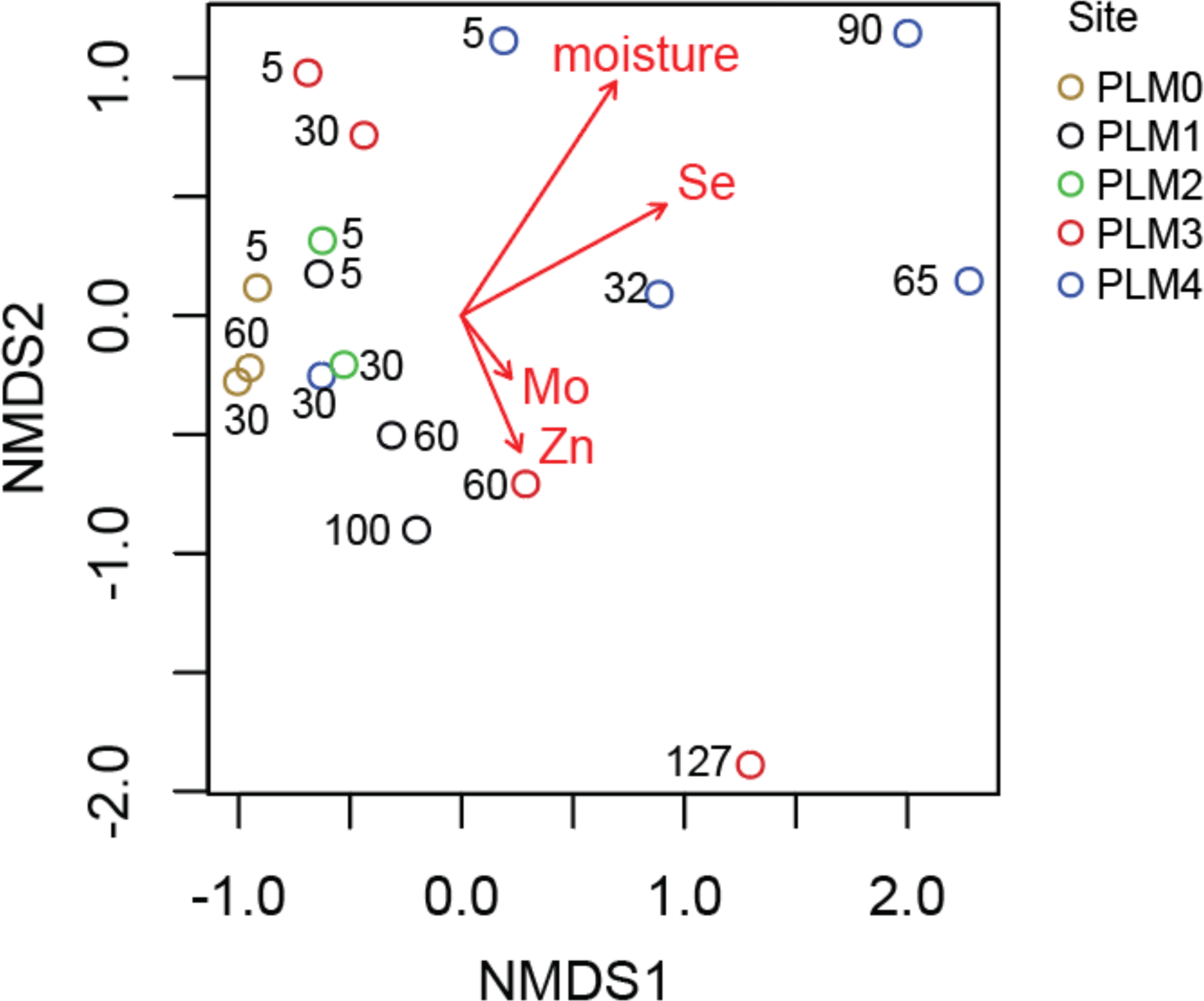
NMDS ordination of microbial communities and correlated geochemical factors. Spearman correlation was tested using Bray-Curtis distances and Euclidean distance matrix. Out of 14 geochemical measurements (Soil moisture, pH, IC, OC, TN, Fe, Se, As, Zn, Mn, Ba, Mo, nitrate, and sulfate) only soil moisture, Se, Mo, and Zn, were correlated with microbial community composition (rho=809). Stress = 0.076. Numbers in figure are depth in cm. Raw values are provided in Table S4.

Metabolic potential, as depicted by detected genes, differentiates locations along the hillslope to floodplain transect. Out of 118 Hidden Markov Models (HMMs), 101 were found to exceed our detection threshold (see Methods). A Principal Component Analysis (PCoA) of gene abundances reveals a clear depth gradient in samples taken from the floodplain site (Fig. 7). A depth-dependent trend in overall metabolic potential is also observed along the hillslope. In addition, gradient in overall metabolic potential correlates with elevation (i.e., position on the hillslope).

**Figure 7:**
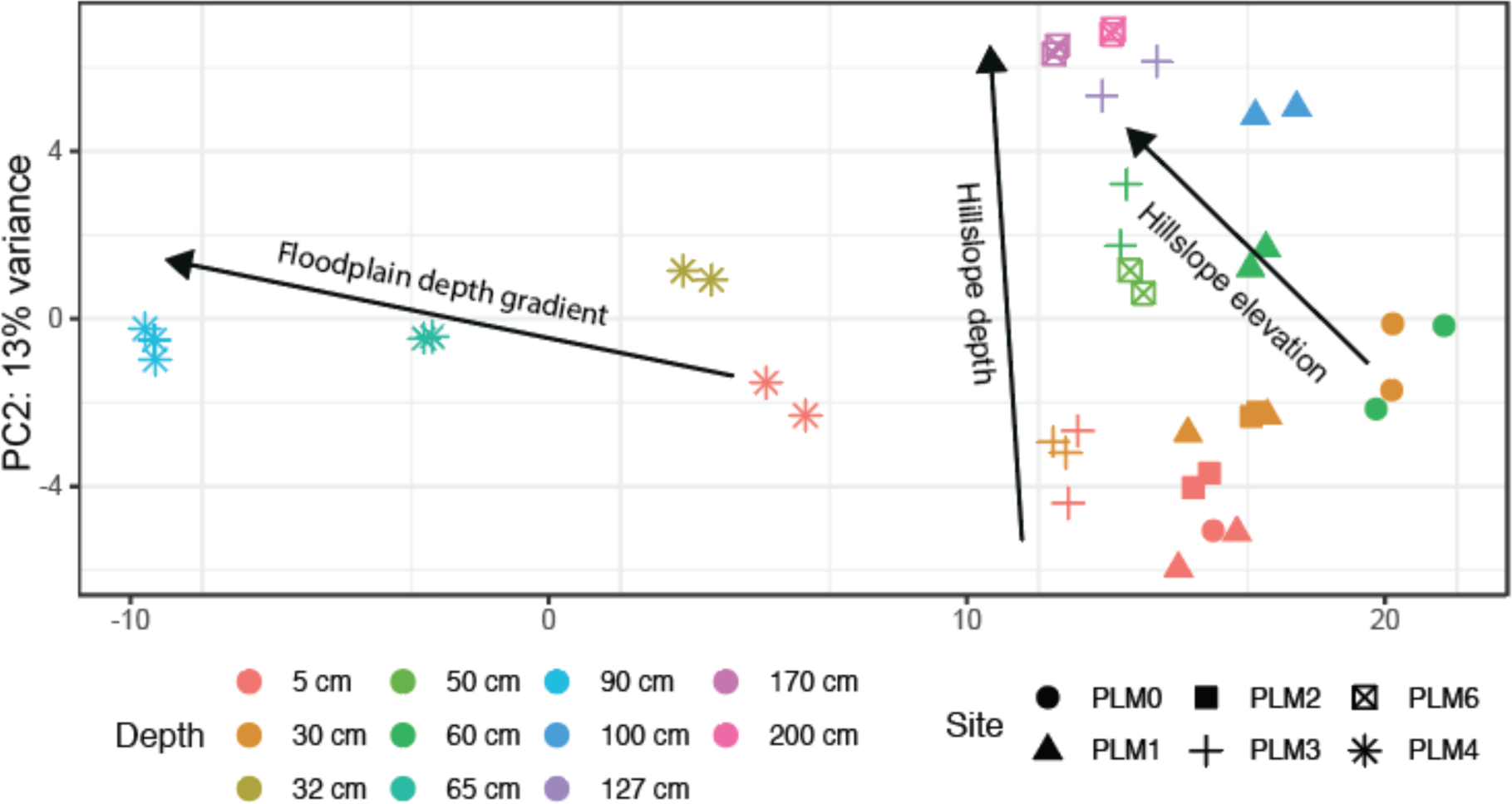
Principal Component Analysis of samples according to abundance of key metabolic enzymes. A. PCoA of key metabolic genes generated using 101 HMMs of carbon, nitrogen, sulfur and selenium metabolic enzymes.

The patterns identified in the PCoA are driven in part by genes encoding enzymes involved in N_2_ fixation, the Wood-Ljungdahl carbon fixation pathway (*cooS*, *cdhC* and *cdhD*) and dissimilatory sulfite reductase subunits A and B (*dsrA* and *dsrB*). All are enriched at the floodplain relative to the hillslope sites, and below compared to above the water table (Fig. 8). Also enriched in samples from below the water table is the catalytic subunit of thiosulfate reductase *phsA,* which catalyzes the reduction of thiosulfate to sulfite and hydrogen sulfide, and [NiFe]-hydrogenases from groups 1, 2a, 2b, 3a, 3b, 3c and 3d. Out of the four genes involved in selenate reduction, *srdA* which is associated with selenate respiration is highly enriched in samples from below compared to above the water table and weathered shale compared to soil.

**Figure 8:**
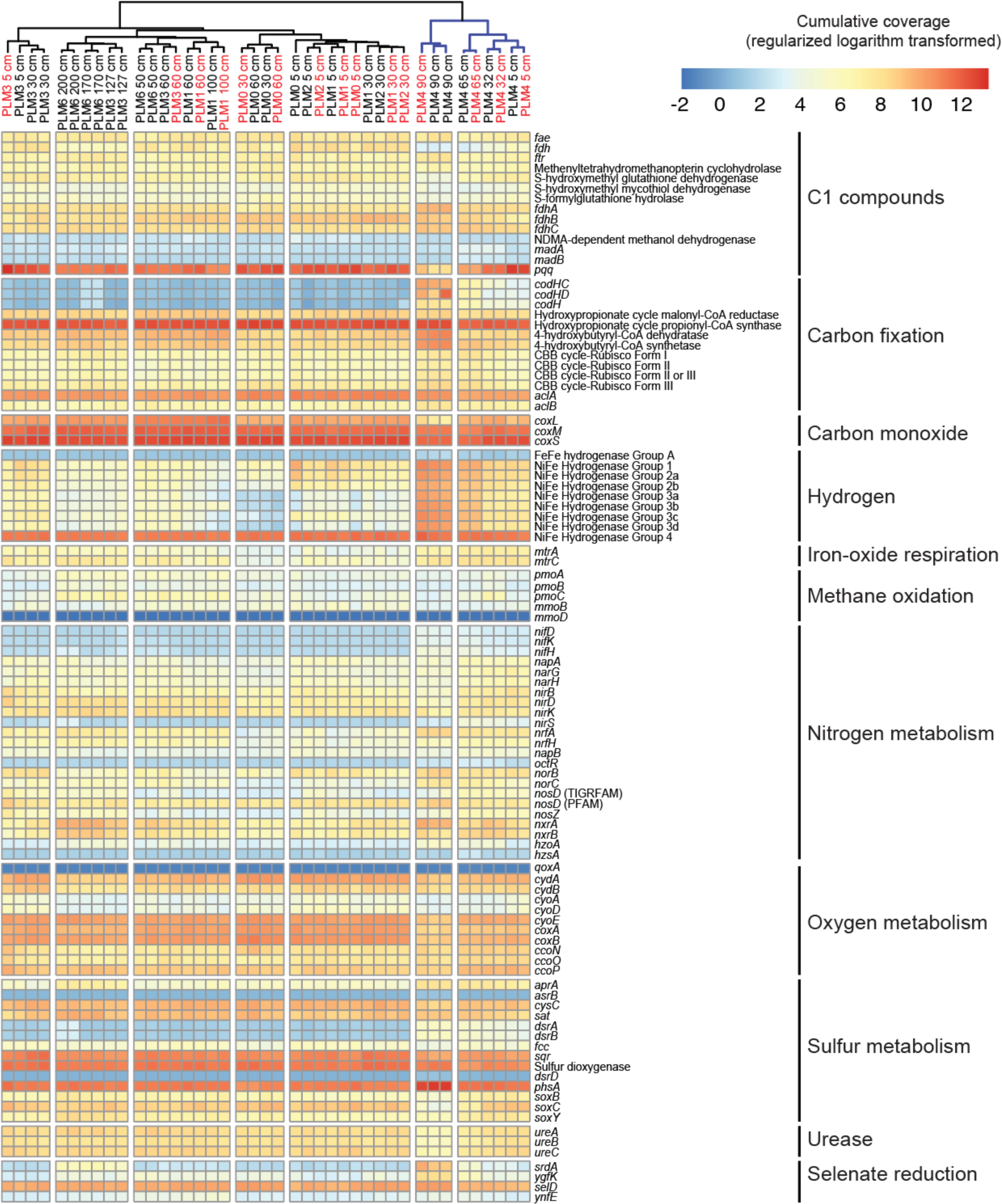
Spatial abundance of genes central to metabolic pathways. Samples from the floodplain (blue colored clade) are distinct from samples of the hillslope (black colored clade), particularly with respect to carbon fixation and selenate reduction. Furthermore, weathered shale samples at PLM6 are distinct from other hillslope samples. Samples names in red denote DNA samples that were co-extracted with RNA (see Methods). The sources of HMMs and their description are given in Table S1.

## Discussion

We integrated metagenomics and soil chemical analysis to investigate how microbial community structure and metabolic potential vary within the subsurface across a transect from high on the East River hillslope to its adjoining floodplain. Our analyses indicate that communities are differentiated according to depth and proximity to weathered shale and groundwater, and that microbial communities of the floodplain soils and sediments differ substantially from those collected along the hillslope.

Notably, the abundance of species of Archaea, Proteobacteria and Candidate Phyla Radiation (CPR) bacteria have distinct spatial patterns. Thaumarchaeota, the dominant archaeal taxon in soils (Bates *et al.*, 2011), are typically aerobic ammonium oxidizers that can drive nitrification (Colman, 2017). They were detected at every depth sampled across the hillslope, as found in hillslope soil pits in Colorado by Eilers et al. (2012). The absence of Thaumarchaeota at the floodplain may be explained by extended periods of water saturation. Low redox conditions, inferred based on abundant genes involved in sulfate and selenate reduction, apparently selected instead for Bathyarchaeota and Euryarchaeota.

Bacteria from candidate phyla increase in abundance with depth throughout PLM sites and may have eluded prior cultivation studies due to their low abundance in more commonly sampled shallow soil. However, CPR bacteria, which elude most cultivation efforts (Solden *et al.*, 2016), are likely dependent on other microorganisms for basic cellular building blocks (Kantor *et al.*, 2013; Brown *et al.*, 2015). Other than the two occurrences of Yanofskybacteria species in deep samples close to the soil-weathered shale transition (127 cm and 170 cm from PLM3 and PLM6, respectively), bacteria from all eight CPR phyla were detected only in the floodplain samples. CPR bacteria are often found in anaerobic environments and have streamlined genomes, lacking many genes for independent survival. Many are likely obligate symbionts, and as such they likely associate with anaerobic hosts with the identity of their hosts remaining unclear (Brown *et al.*, 2015; Hug, Baker, *et al.*, 2016; Castelle and Banfield, 2018).

The abundance of genes encoding methanol dehydrogenase (pqq in Figure 8) and the catalytic subunit of carbon monoxide dehydrogenase (coxL in Figure 8) were consistently lower in the groundwater-saturated floodplain samples than in any hillslope samples or floodplain samples from above the water table. Methanol dehydrogenase is involved in aerobic oxidation of methanol (which could derive from plant biomass or oxidation of methane), whereas CO dehydrogenase is involved in aerobic oxidation of CO (possibly produced by plants as a signaling molecule). Sulfate reduction may be a second biogeochemical process that differentiates microbial communities at the floodplain from those on the hillslope, particularly in samples below the water table where genes involved in this process are relatively abundant. Further, three genes encoding for key enzymes in the anaerobic Wood-Ljungdahl pathway for carbon fixation and genes for nitrogen fixation are relatively abundant at the floodplain site compared to the hillslope sites. These patterns support the conclusion that groundwater-saturated regions of the watershed support largely anaerobic microbial communities. Overall, the findings indicate that floodplain site metabolic potential is depth-stratified, with one microhabitat below the water table and apparently colonized by organisms with anaerobic metabolisms and another above the water table where communities would experience fluctuating states of aeration. The spatial layout of the two compartments may support complete redox cycles, analogous to sulfur cycling at oxygen-minimum zones in the ocean (Canfield *et al.*, 2010).

Selenium concentrations may be a major factor that differentiates microbial communities at the floodplain from those on the hillslope. Selenium occurs in insoluble metal selenides in Mancos Shale that underlies much of the Gunnison River basin (Colorado, USA; Elrashidi, 2018), which includes the East River watershed. Oxidation of selenium to soluble selenite and selenate under mildly reducing to oxidizing conditions (Presser, 1994) leads to its mobilization and probably accounts for its presence in pore fluids. Enrichment of *srdA* genes, which encode the catalytic subunit of the complex required for selenate reduction in sequences from the floodplain site suggests that dissimilatory reduction of selenate (Maiers *et al.*, 1988; Ike *et al.*, 2000; Williams *et al.*, 2013; Fakra *et al.*, 2015; Nancharaiah and Lens, 2015) supports microbial growth at this site. *Geobacter* species, which were identified almost exclusively in floodplain samples (Fig. S4, clade 3) and are sometimes capable of selenite reduction (Pearce *et al.*, 2009), may be responsible for these reactions. The detection of *srdA* genes in the three deepest samples from the hillslope suggests that selenate reduction may occur periodically close to the weathered shale-soil interface where seasonally variable redox conditions induced by groundwater fluctuations may enable microbe-catalyzed selenium transformations.

It is interesting to note that across the hillslope sites, shallow soils have relatively similar community compositions. This might be explained by the low soil moisture that these locations experience over much of the year and exposure to low temperatures during late fall and early winter prior to the onset of insulating snow cover. Further, soil community compositions are homogenized at some sites, likely due to soil mixing as a result of gopher activity (Yoo *et al.*, 2005) that may also increase soil porosity and permeability and homogenize the mineral matrix and microbial community composition within a site, particularly close to the soil surface (reviewed by Platt *et al.*, 2016). It is also possible that similarity in vegetation at the non-floodplain sites contributes to community similarity.

Several factors may contribute to variation in microbial community composition across the sites that cannot be explained by observed physiochemical differences. Between site heterogeneity, possibly due to periodic events or local changes in vegetation, could be eliminated by microbial dispersal. However, microbial dispersal is generally very limited in soils that are not saturated with groundwater (Elsas *et al.*, 1991). Although groundwater and runoff from rain and snowmelt might transport microbes down slope and into the weathered rock, hydraulic measurements show that overland and lateral underground transport is likely limited at the hillslope sites (Tokunaga et al. in prep). Soil and weathered rock are water saturated for only a few weeks each year other than at the floodplain. During this period, water moves at ∼10 to 20 m per month parallel to the surface slope (Tokunaga et al. in prep), distances that are too short to connect communities at our sampling sites.

Our study of a hillslope lower montane meadow to floodplain transect revealed an ecosystem comprised of distinct subsystems. Specifically, our results documenting the abundance patterns of genes involved in selenium, sulfur, carbon and nitrogen cycles suggest that hillslope and floodplain sites constitute distinct microbial habitats. Further, the hillslope sites are spatially differentiated into microhabitats close to (or within) weathered shale and proximal to the surface. Similarly, the floodplain site is resolved into largely anaerobic and aerobic communities over relatively short vertical distance, raising the possibility of elemental cycling across the interface. These results clarify the scale of heterogeneity in biogeochemical processes and improve our understanding of how these processes map onto the watershed.

## Acknowledgments

**Yongman Kim** – collecting samples for chemistry, and soil chemistry analysis.

**Wendy Brown** – Information about gopher activity and vegetation at PLM sites

The work described in the manuscript was supported as part of the Watershed Function Scientific Focus Area funded by the U.S. Department of Energy, Office of Science, Office of Biological and Environmental Research under Award Number DE-AC02-05CH11231.

## Supplemental Figures

**Figure S1: Illustration of the East River watershed**. Star and flag indicate the town of Crested-Butte and the location of the Pumphouse Lower Montane (PLM) sampling site, respectively.

**Figure S2: Soil texture of samples from PLM0, PLM1, and PLM3.** All soils are categorized as silty-loam (A) with the greatest variability being 20-60% sand content. PLM0 and PLM1 are found to contain more sand than in PLM3 (B) and sand content is correlated with the PC1 which explains almost 80% of the variability between the samples.

**Figure S3: Geochemistry measurements of elements that showed correlation to microbial community structure.** Data is missing for PLM6. Sulfate measurements are missing for PLM0 40 cm, PLM1 60 cm and 90 cm, PLM2 5 cm and 30 cm.

**Figure S4: A Maximum-Likelihood phylogenetic tree of rpS3 clusters classified as Deltaproteobacteria.** Black circles mark branch support greater than 0.8. Grey scale bar was calculated with the square root of relative abundance of each cluster. Clades of interest are marked 1 through 5. The phylogenetic tree is provided as Supplemental File 1.

## Supplemental Tables and Files

**Table S1 - Key genes in metabolic pathways and the origin of the HMM model that was used to identify them in the metagenomic samples.**

**Table S2 - Sequencing depth and assembly information**

**Table S3 - Soil texture**

**Table S4 - Soil chemistry data**. Ratio of 1:1 (soil:DIW mass ratio) extraction. The original porewater was considered as a part of the total water mass.

**File1 – Hidden Markov Model of *srdA*.**

**File2 – phylogenetic tree of rpS3 amino-acid sequences in Newick format.**

